# Exploiting codon usage identifies RpS21 as an *in vivo* signal strength-dependent Ras/MAPK regulator

**DOI:** 10.1101/650630

**Authors:** Jessica K. Sawyer, Zahra Kabiri, Ruth A. Montague, Sarah V. Paramore, Erez Cohen, Hamed Zaribafzadeh, Christopher M. Counter, Donald T. Fox

**Affiliations:** Department of Pharmacology & Cancer Biology, Duke University Medical Center, DUMC Box 3813, Durham, NC 27710; Department of Cell Biology, Duke University Medical Center, DUMC Box 3813, Durham, NC 27710; Duke Cancer Institute, Duke University Medical Center, DUMC Box 3813, Durham, NC 27710

**Keywords:** *Drosophila*, Ras, codon bias, RpS21

## Abstract

Signal transduction pathways are intricately fine-tuned to accomplish diverse biological processes. An example is the conserved Ras/mitogen-activated-protein-kinase (MAPK) pathway, which exhibits context-dependent signaling output dynamics and regulation. Here, by altering codon usage as a novel platform to control signaling output, we screened the *Drosophila* genome for modifiers specific to either weak or strong Ras-driven eye phenotypes. We mapped the underlying gene from one modifier to the ribosomal gene *RpS21*. RpS21 preferentially influences weak Ras/MAPK signaling outputs, and negatively regulates Ras/MAPK in multiple cell/tissue and signaling settings. In turn, MAPK signaling may regulate its own negative feedback by promoting RpS21 expression. These data show that codon usage manipulation can identify output-specific signaling regulators, and identify RpS21 as an *in vivo* Ras/MAPK phenotypic regulator.

## INTRODUCTION

Conserved signal transduction pathways are employed reiteratively throughout nature during diverse processes such as cell fate decisions and tissue growth. These same pathways can be aberrantly regulated in disease. Large numbers of molecular regulators of these pathways have been identified using high-throughput genetic screening. Additionally, quantitative imaging approaches have revealed intricate signaling regulation. This regulation includes feedback control of the duration or strength of a downstream biochemical signaling output (e.g., weak or strong activation of a target gene). A current challenge is to place the numerous identified signaling pathway regulators in the context of complex signaling dynamics, and to relate such regulation to *in vivo* signal-dependent processes.

An example of the complexity of signaling regulation is the evolutionarily conserved Ras/mitogen activated protein kinase (MAPK) pathway. In canonical MAPK signaling, receptor tyrosine kinase stimulation converts the Ras GTPase to an active GTP-bound conformation. Ras-GTP then activates the MAPK pathway, comprised of Raf kinases, which are activated by Ras and phosphorylate/active Mek kinases, which do the same to Erk kinases. Through highly successful modifier screen approaches in models such as the *Drosophila* eye^1–8^ and *C. elegans* vulva^9–13^, regulators of this core pathway have been identified. Additional *Drosophila* cell-based screens using a biochemical MAPK output (Erk phosphorylation) have identified many other Ras/MAPK regulators^14–16^.

These numerous molecular regulators contribute to a diversity in Ras/MAPK signaling dynamics. Using an optogenetics-driven MAPK activation approach in cultured mammalian cells, it was revealed that distinct Ras/MAPK regulation (such as a paracrine STAT3 circuit) can distinguish between biochemical signaling outputs, namely sustained (strong) or transient (weak) Erk activation by Ras^17^. These biochemical outputs are regulated by negative feedback on Erk^18^.

Importantly, *in vivo* context plays a role in whether a given strength of signaling output leads to a phenotypic output. Specifically, taking a similar optogenetic approach in the early developing fly embryo, it was shown that manipulating Erk activation strength has minimal effects on cells at the poles of the embryo, but has a profound impact on development of cells in the middle of the embryo^19^. Further, expressing an activating mutant of Mek in either *Drosophila* or zebrafish was recently shown to either activate or repress Erk phosphorylation depending on the cell type and gene expression environment^20^. These observations suggest that distinct Ras/MAPK regulation operates in distinct cellular contexts. Taken together, these studies highlight the need to better understand how distinct Ras/MAPK signaling states (e.g., strong or weak) are controlled by distinct sets of Ras/MAPK molecular regulators, in the context of an *in vivo* phenotype.

Here, we introduce an approach to genetically screen for signal output-specific regulators of Ras/MAPK signaling. This approach, which should be applicable to any signaling pathway, involves controlling the amount of active Ras protein produced by changing codon usage in the single *Drosophila Ras* gene^21^ (FlyBase: *Ras85D,* hereafter *Ras)*. Rare codons are well-associated with poor mRNA translation^22^. We have previously found that changing rare codons in the mammalian Ras isoform *KRAS* to their common counterparts leads to elevated translation, protein, signaling, and transformation^23^. We report the generation and characterization of transgenic flies and cell lines whereby the amount of active Ras protein produced, the resultant level of Erk activation, and resultant rough-eye phenotype is dictated solely by the codon usage engineered into a given *Ras* transgene. We then report the use of such transgenic flies to screen a whole genome deficiency (termed *Df* for convenience) kit for genetic modifiers of eye phenotypes that are specific to only strong or only weak Ras/MAPK signaling. Of the 15 *Dfs* identified, we successfully mapped the modification of *Df(2L)BSC692*, an enhancer of the rough-eye phenotype driven only by weak Ras/MAPK signaling, to the ribosomal protein S21 gene (*RpS21*). We show that RpS21 negatively regulates Ras protein levels in several contexts, the effect of which is preferentially manifested at low levels of MAPK signaling. Further, MAPK signaling may also positively regulate RpS21 protein levels, suggesting that RpS21 is potentially part of a negative feedback regulatory mechanism that is most effective under conditions of low-level MAPK signaling. This approach highlights the usefulness of codon manipulation as a viable approach to identify signal output-specific signaling regulation.

## RESULTS

### Exploiting codon usage to control MAPK signaling output

To identify Ras/MAPK molecular regulators that differentially impact strong or weak signaling outputs, we required a platform to tightly control the strength of MAPK signaling. To activate the pathway, we expressed a highly conserved, mutant active (G12V) *Drosophila Ras* transgene (termed *Ras*^V12^ here for convenience). To control MAPK signaling strength during fly development, we opted for the new approach of simply changing the codon usage of a *Ras^V12^* transgene. Codons that occur infrequently in a given genome (rare codons) are known to impede translation, including in *Drosophila*^24–30^. By engineering a gene enriched in rare codons for each given amino acid, it is possible to create an mRNA that is poorly translated without altering the amino acid sequence of the encoded protein. This has the distinct advantage that control of protein expression is embedded in the DNA and requires no additional factors or experimental variables. We used established data on *Drosophila* codon usage (see Methods) and created four distinct versions of *Drosophila Ras* transgenes: *1*) we altered none of the codons (*Ras^V12^Native*), *2*) we made all codons the most commonly occurring in the genome (*Ras^V12^Common*), *3*) we made all codons the most rare in the genome (*Ras^V12^Rare*), and *4*) we created a control wild-type version lacking the V12 mutation and also lacking codon alteration (*Ras^WT^Native*). To monitor expression, all four transgenes were epitope-tagged at the N-terminus with a 3XFLAG-tag sequence and expressed under the control of a Gal4-inducible UAS promoter (Fig 1a, see Methods). We note that *Ras^V12^Native* has primarily common codons and a similar Codon Adaptive Index (CAI^31^) to *Ras^V12^Common*^21^, while the CAI for *Ras^V12^Rare* is much lower (**FigS1a, b**). To control for position effects, all transgenes were integrated at the same site in the genome (see Methods).

**Figure 1.**
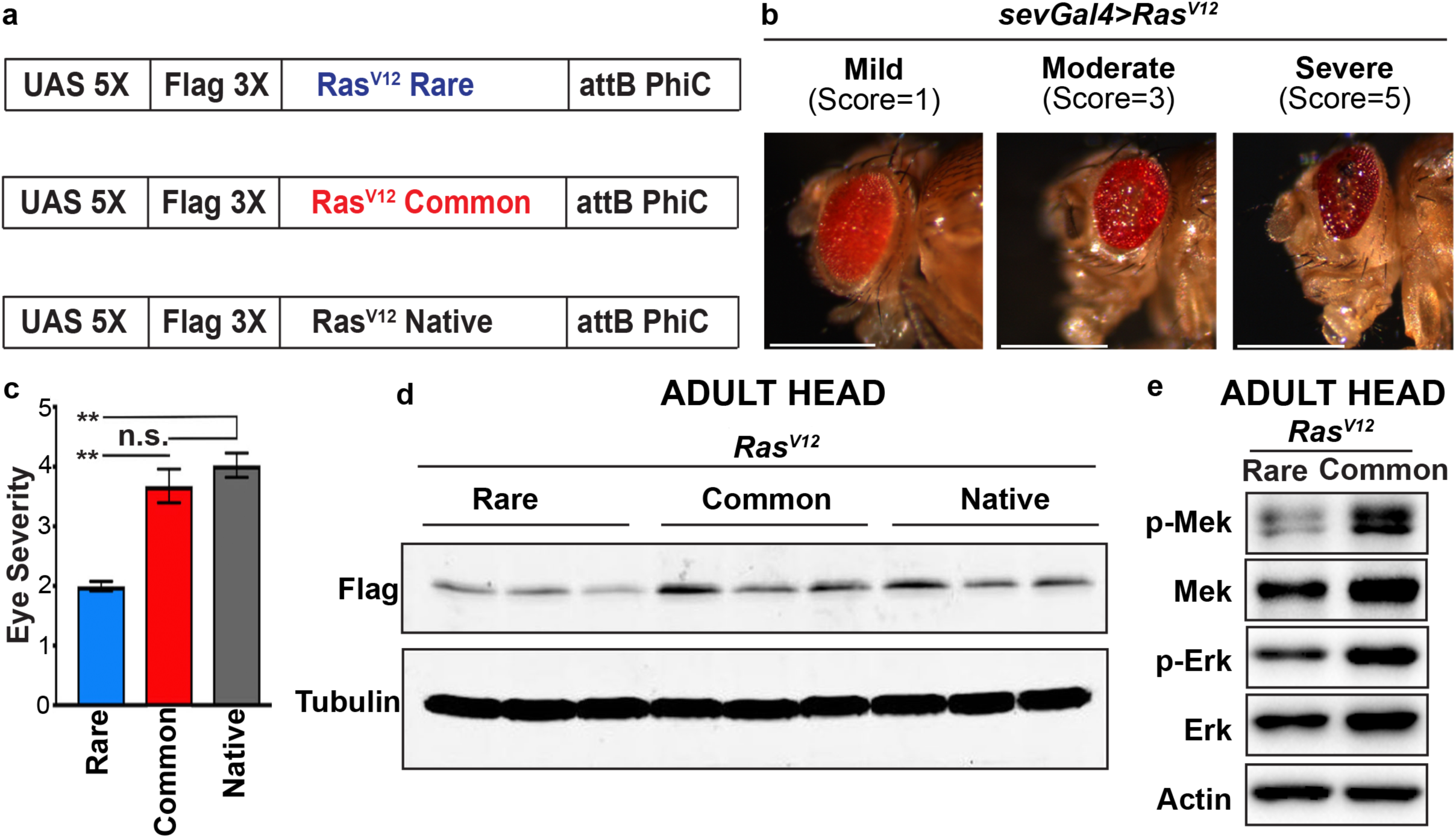
Exploiting codon usage to control MAPK signaling output. **(a)** Schematic representation of the FLAG epitope-tagged *Ras^V12^*transgenes encoded by rare, common, or native codons. **(b)** Images representing the eye phenotypes assessed and the scoring system. Scale bars = 0.5mm. **(c)** The mean ± SEM eye severity score of the indicated *Ras^V12^* transgenes from three replicate experiments at 25°C. Tukey’s multiple comparisons test was used for statistical comparisons. \*\*\*\**p*<0.0001. \*\*\**p*<0.001. \*\**p*<0.01. n.s., not significant. **(d)** Immunoblot detection transgenic Ras^V12^ protein (with an anti-FLAG antibody) and αTubulin as a loading control from lysates derived from the head of flies with the indicated versions of transgenic *Ras^V12^*. **(e)** Immunoblot detection of phosphorylated (p-) and total Mek and Erk, and actin as a loading control from lysates derived from the head of flies with the indicated versions of transgenic *Ras^V12^*.

As a measure of signaling output strength, we chose to use an *in vivo* phenotypic readout rather than a biochemical readout, an approach validated by quantitative studies of MAPK activation in *Drosophila* embryos^19, 20^. For genetic screening of Ras/MAPK phenotypic regulators, the *Drosophila* eye is a highly accessible model. Driving expression of *Ras^V12^* in the developing eye with an eye-specific promoter such as *sevenless* (*sev*) dysregulates the proper differentiation of the R7 photoreceptor cell, leading to an easily scored ‘rough-eye’ phenotype^32, 33^. This phenotype relies on Ras action through the conserved MAPK pathway^1, 34^.

We first assayed the phenotypic output of each *Ras* transgene *in vivo* by driving their expression in the developing fly eye using *sevenless (sev)-Gal4*. As expected^32^, expression of *Ras^WT^* in this manner does not result in a rough-eye phenotype (**FigS1c**). However, when we expressed the constitutive-active versions of *Ras,* we found a range of rough-eye phenotypes (Fig 1b). We binned these phenotypes into one of three classes: severe, moderate, or mild. Each class was assigned an increasing numeric score (Fig 1b, see Methods). We then calculated an average severity score for each *Ras* transgene. *Ras^V12^Native* and *Ras^V12^Common* animals exhibit a similar phenotypic score, reflecting their similar CAI. Further, this phenotypic score is, on average, approximately 2-fold more severe than that of *Ras^V12^Rare* (Fig 1c). To determine whether protein levels track with the difference in rough-eye phenotype, we isolated heads from flies encoding the three active *Ras* transgenes in triplicate and immunoblotted with an anti-FLAG antibody. Ras^V12^ protein levels are similar between *Ras^V12^Native* and *Ras^V12^Common* flies, and both are expressed ∼2-fold higher than *Ras^V12^Rare* (Fig 1d, **FigS1d**). Given that *Ras^V12^Native* and *Ras^V12^Common* produce similar levels of protein and have the same rough-eye phenotype, for further experiments we opted to control *Ras^V12^* expression using the *Ras^V12^Common* and *Ras^V12^Rare* transgenes. These experiments established that codon usage can be manipulated to examine an *in vivo, Ras* signal-driven output (eye phenotype), and identified both weak (*Ras^V12^Rare*) and strong (*Ras^V12^Common*) versions of this output.

To examine the effect of expression of *Ras^V12^Rare* versus *Ras^V12^Common* transgenes on known biochemical outputs of downstream MAPK signaling, we evaluated the level of activated MAPK signaling in fly heads by measuring the level of phosphorylated Mek (p-Mek, FlyBase: *Dsor*) and Erk (p-Erk, FlyBase: *rolled*) compared to the total level of these proteins by immunoblot analysis. Compared to *Ras^V12^Rare* animals, *Ras^V12^Common* animals exhibit elevated levels of p- Erk (and to a lesser extent p-MEK) compared to *Ras^V12^Rare* fly heads (Fig 1e**, FigS1e**). We independently verified this difference in cultured S2 and KC insect cells (see Methods), again finding that *Ras^V12^Common* is expressed higher and more robustly activates the MAPK pathway compared to *Ras^V12^Rare* (**FigS1f**). In sum, our findings establish *Ras^V12^Rare* and *Ras^V12^Common* as two distinct transgenes that either weakly or strongly control Ras/MAPK signaling output (as measured by *Ras^V12^*expression and MAPK activation), and that transgene-driven signal strength tracks with an observable difference in phenotypic output.

### A genome-wide screen uncovers differential phenotypic regulation between strong and weak Ras/MAPK signaling states

We next sought to use our codon alteration system to gain insight into how the Ras/MAPK pathway can be differentially regulated in different signal-strength states. To do so, we screened for molecular regulators that modify Ras/MAPK phenotypes driven only by strong or only by weak signaling states. We first confirmed that *Ras^V12^Common* and *Ras^V12^Rare* rough-eye phenotypes were both in the range that can be modified. Specifically, heterozygous loss-of-function mutations in two known *Ras* suppressors, ***<u>k</u>****inase **<u>s</u>**uppressor of **<u>r</u>**as,* or *ksr*^8^, and ***<u>b</u>****eta subunit of type I **<u>g</u>**eranyl***g***eranyl* t*ransferase*, or *betaggt-I*^2^, suppress the rough-eye phenotype for both *Ras^V12^Common* and *Ras^V12^Rare* (**FigS2a**). Conversely, we examined heterozygous mutants of the known *Ras* enhancer ***<u>a</u>****nterior **<u>op</u>**en*, or *aop*^35^. Although we did not observe clear eye enhancement, we did observe a marked decrease in survival of *aop/+* animals expressing both *Ras^V12^Common* and *Ras^V12^Rare* transgenes (**FigS2b**). This result likely reflects the expression of *sevenless-Gal4* in other tissues^36^. These results establish that codon-altered *Ras^V12^* transgenes are subject to phenotypic modification, including by dose-sensitive haplo-insufficient mutations.

Previous modifier screens, including in the eye, employed the native *Ras* cDNA to express activated *Ras*^1,7, 33, 37^. This sequence has a strong common-codon bias (**FigS1b**), and is similar to *Ras^V12^Common* in terms of MAPK biochemical and phenotypic outputs (Fig 1). To find unidentified modifiers that may be specific to weaker (or stronger) MAPK-driven phenotypes, we conducted a genome-wide unbiased haplo-insufficiency screen to specifically identify modifiers of the rough-eye phenotype driven by only *Ras^V12^Rare,* (or only *Ras^V12^Common*), (Fig 2a). We used the Bloomington Deficiency (*Df*) Kit, which covers 98.3% of the euchromatic genome^38^. In a primary screen (Fig 2b**, TableS1**), we crossed 470 *Dfs* representing 99.1% of the *Df* collection to animals with *Ras^V12^Rare* or *Ras^V12^Common* expressed in the eye by *sev-Gal4*, and scored the resulting eye severity in an average of 30 (*Ras^V12^Common*) or 60 (*Ras^V12^Rare*) progeny animals per cross. We also factored animal lethality into our scoring (see Methods).

**Figure 2.**
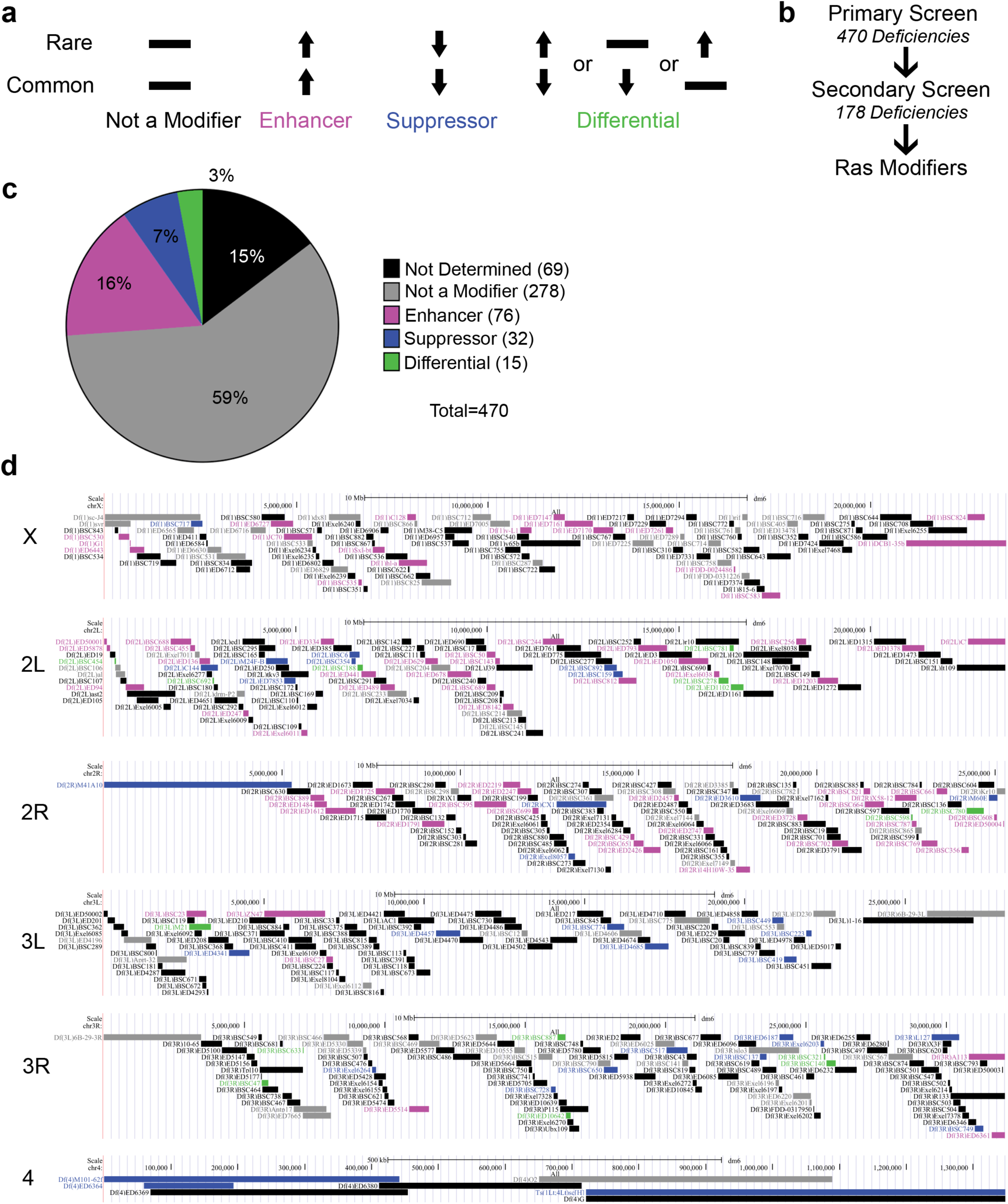
A genome-wide screen uncovers differential phenotypic regulation between strong and weak Ras/MAPK signaling states. **(a)** Schematic of the *Ras* modifiers types scored in the *Df* screen. **(b)** Schematic of screening approach. **(c)** Pie chart showing the number of *Df* with the indicated types of *Ras* modifiers. **(d)** Genome map of deficiencies color coded as in **c** for the class of *Ras* modifier.

As expected, we found general *Ras* modifiers that either enhance or suppress eye phenotypes driven by both *Ras^V12^* transgenes (Fig 2c, d**, TableS1**). Interestingly, we identified more enhancers than suppressors (16% versus 7%, Fig 2c). The reason for this remains to be determined, but we note that our calculation of phenotypic modification (see Methods) included scoring animal lethality, which may identify strong enhancers of *Ras^V12^Common* not identified in previous screens based solely on a rough-eye phenotype. Of great interest, we also identified *Dfs* whereby *Ras^V12^Common* and *Ras^V12^Rare* are differentially modified (Fig 2a), meaning they scored as only modifying the eye phenotype driven by a single signaling state (*Ras^V12^Common* or *Ras^V12^Rare,* not both). Using a low-stringency cutoff score (see Methods), we identified 178 putative differential modifier *Dfs* in our primary screen (Fig 2b**, TableS1**). These *Dfs* were then re-tested in a secondary screen (Fig 2b) by crossing them a second time to *sev-Ras^V12^Common* and *sev- Ras^V12^Rare*. In this screen, we used a more stringent cutoff score to define a differential modifier (see Methods). This scoring and replicate analysis reduced the number of candidates to 15 *Dfs*, or 3% of the tested *Dfs* (Fig 2c, d**, TableS1**), that reproducibly differentially modify either only *Ras^V12^Common* or only *Ras^V12^Rare* (Fig 3a). Among these differential modifiers, we again recovered more enhancers than suppressors, although importantly we recovered both enhancers of *Ras^V12^Common* and suppressors of *Ras^V12^Rare*, arguing that our screen had the dynamic range to modify both strong (*Ras^V12^Common*) and weak (*Ras^V12^Rare*) Ras/MAPK signaling outputs (Fig 3b).

**Figure 3.**
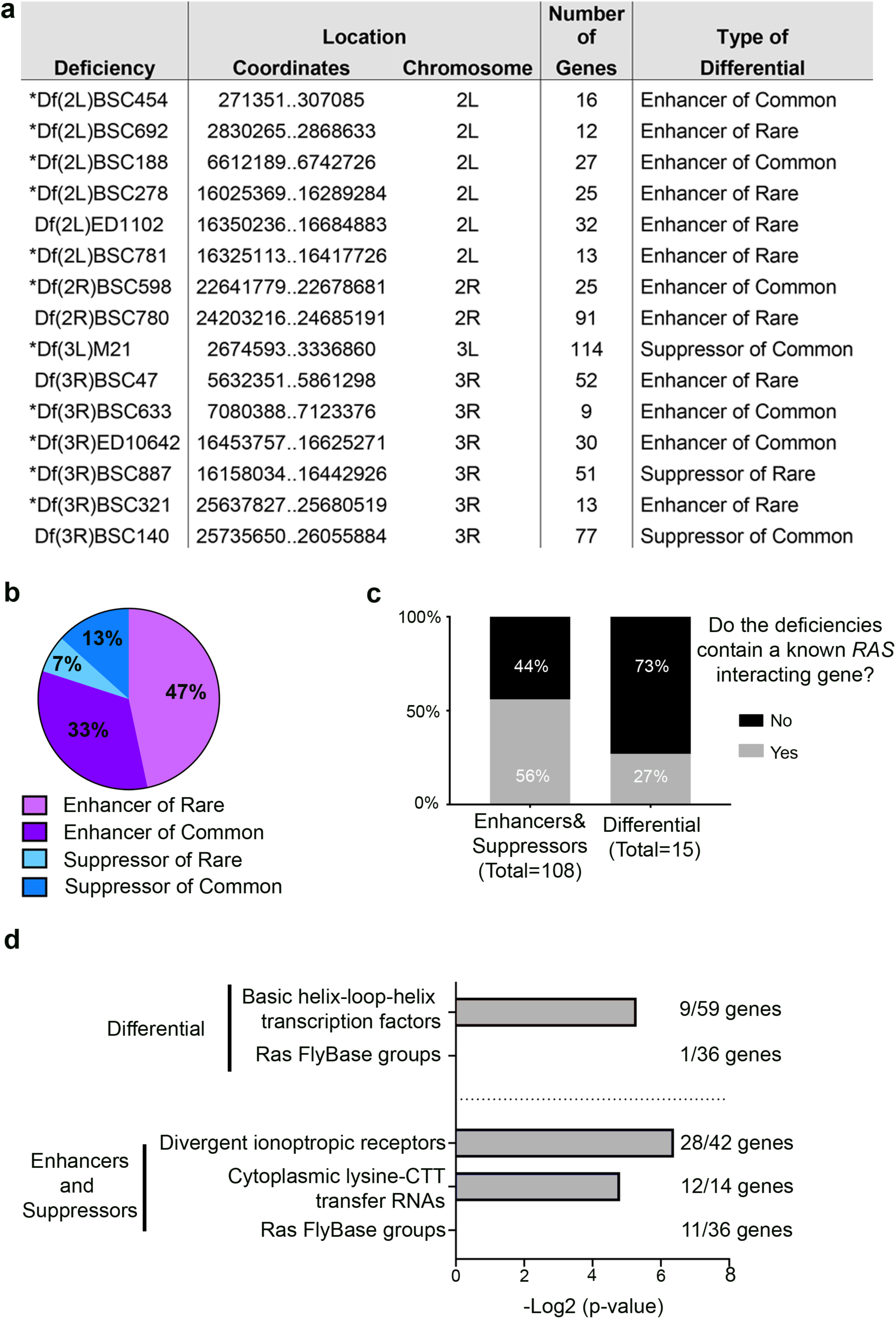
Characterization of differential modifiers. **(a)** Characterization of the differential *Ras* modifiers identified. Asterisks= those *Df*s for which no known *Ras* modifier has been reported (see Methods). **(b)** Pie chart showing the number of differential modifiers with the indicated phenotypes. **(c)** Graph of the percent (and number) of *Dfs* that do or do not contain known *Ras* interacting genes. **(d)** Enriched FlyBase gene groups contained in differential versus enhancer and suppressor deficiencies.

We next queried both the general (signal output-independent) and differential (signal output-dependent) modifiers against a FlyBase database of all reported *Ras* genetic enhancers and suppressors (see Methods). 56% of our general modifier *Dfs* covered regions of the genome containing reported *Ras* enhancers or suppressors, validating our approach. Additionally, we note that among our identified differential modifier *Dfs*, most (73%) do not encompass known *Ras* modifiers, supporting the idea that our signal strength-specific modifier hits are enriched in new *Ras* enhancers and suppressors (Fig 3c). To explore possible relationships amongst these 15 differential modifier *Dfs*, we queried the genes within differential versus enhancer and suppressor *Dfs* against the established list of FlyBase Gene Groups (FBGG). Interestingly, the gene groups enriched in the differential *Dfs* do not overlap with those in the general enhancer/suppressor *Dfs* (Fig 3d), suggesting that the differential modifiers may represent a distinct class of Ras modifiers. Unlike the general modifier Dfs, differential modifier regions are enriched for basic Helix Loop Helix (bHLH) transcription factors, potentially reinforcing their distinct regulation of Ras/MAPK signaling. In summary, by controlling Ras/MAPK signal output strength through codon usage and using a phenotypic output screen, we successfully identified *Dfs* that alter a Ras/MAPK phenotype in a signaling output-specific fashion.

### RpS21 negatively regulates Ras/MAPK signaling in a signal strength-specific manner

To identify a differential modifier at the single gene level, we focused on *Df(2L)*BSC692 as it was one of the smallest deficiencies, encompassing only 12 genes, that specifically enhanced *Ras^V12^Rare* (Fig 3a**, FigS3a**). Of these 12 genes, ***<u>R</u>****ibosomal **<u>p</u>**rotein* **<u>S21</u>**, or RpS21 (also known as *overgrown hematopoietic organs 23B/oho23B)*, represented a plausible candidate modifier. RpS21 stands out among small ribosomal subunits for its reported negative regulation of hematopoietic and imaginal disc hyperplasia^39^. To determine if *RpS21* was the responsible gene in *Df(2L)BSC692* for specifically enhancing *Ras^V12^Rare*, we assessed the rough-eye phenotype of *Ras^V12^Common* and *Ras^V12^Rare* in the background of the mutant *RpS21^0375^*. Indeed, only the *sev-Ras^V12^Rare* rough-eye phenotype is enhanced in the *RpS21^0375^/+* background (Fig 4a). To test whether other ribosomal subunits may also be involved in Ras/MAPK regulation, we conducted a screen using available heterozygous mutants and RNAi constructs for 15 other small ribosomal subunits. Where possible, we used multiple alleles/RNAi constructs. Of these, at least one allele/RNAi for 4 other subunits also scored as differential (signal-strength specific) Ras/MAPK modifiers in the eye, including mutants for 3 genes that, like *RpS21* scored as enhancers of *Ras^V12^Rare* but not *Ras^V12^Common* (**TableS2**). Together, these findings identify *RpS21* as the responsible modifier in one *Df* from our signal strength-specific screen, and also implicate at least a subset of other ribosomal subunits in signal strength-dependent Ras/MAPK regulation.

**Figure 4.**
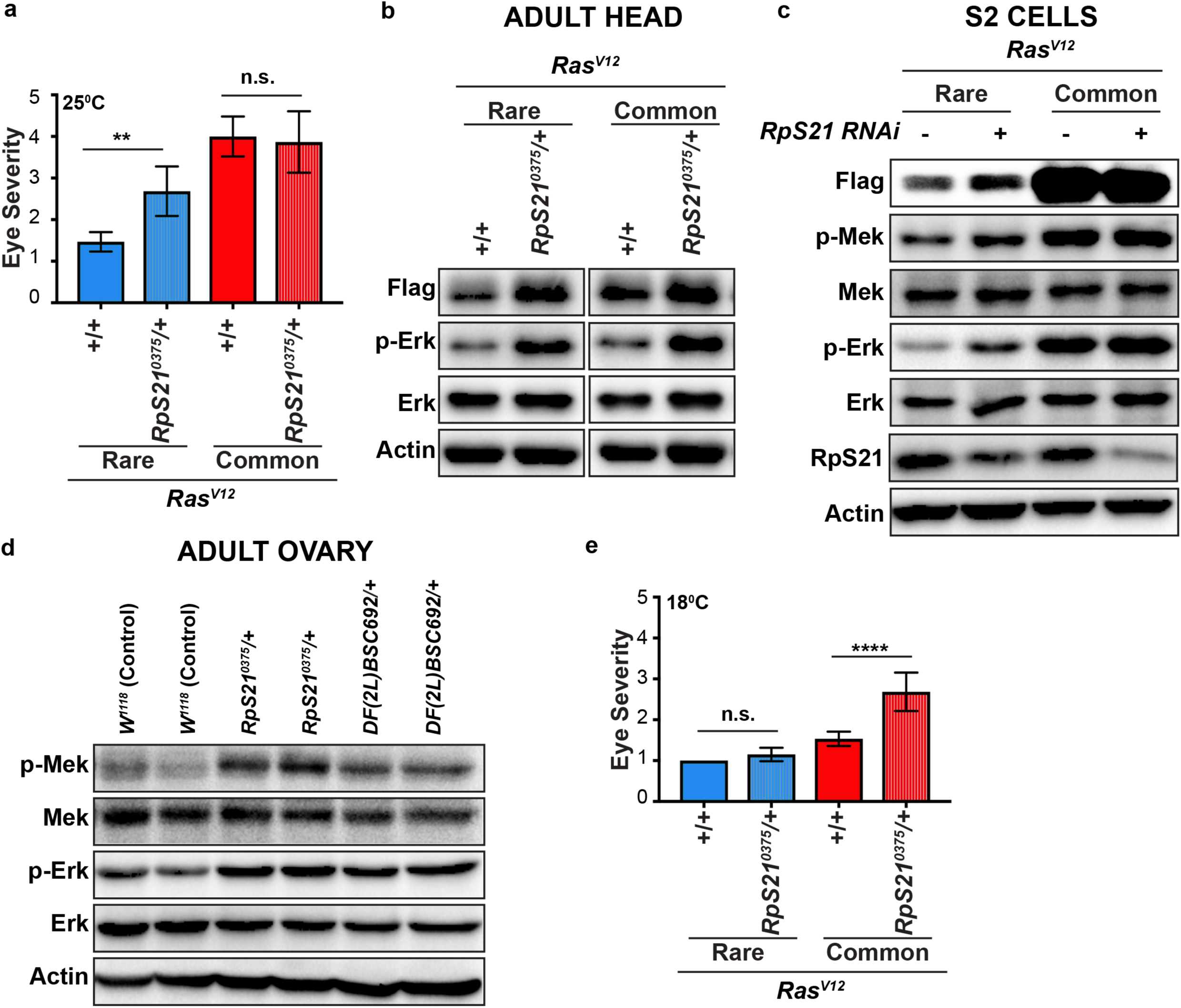
RpS21 negatively regulates Ras/MAPK signaling in a signal strength-specific manner. **(a)** The mean ± SEM eye severity score of the genotypes from three replicate experiments at 25°C. **(b,c,d)** Immunoblot detection of transgenic Ras^V12^ (with an anti-FLAG antibody), phosphorylated (p-) and total Mek and/or Erk, RpS21, and actin as a loading control from lysates derived from **(b)** the heads of flies with the indicated versions of transgenic *Ras^V12^* in either the wild-type (+/+) or mutant (*RpS21^0375^/+)* RpS21 backgrounds, **(c)** S2 cells stably transduced with expression vectors expressing the indicated *Ras^V12^* transgenes in the absence (-) and presence (+) of *RpS21* RNAi, or **(d)** the ovaries of either wild-type (*+/+*) or mutant (*RpS21^0375^/+*) flies. **(e)** The mean ± SEM eye severity score of the genotypes from three replicate experiments at 18°C. Tukey’s multiple comparisons test was used for statistical comparisons in **b** and **e**. \*\*\*\**p*<0.0001. \*\**p*<0.01. n.s., not significant.

Given the identification of both an *RpS21* mutant allele and a small deficiency encompassing this gene (*Df(2L)BSC692)* as a differential *Ras^V12^* modifier, we next turned our attention to the underlying mechanism of this phenotype. To this end, we measured the level of Ras^V12^ and p-Erk by immunoblot analysis in four completely distinct settings: ectopic signaling in adult fly heads, ectopic signaling in cultured S2 cells, endogenous signaling in cultured serum-stimulated KC cells, and endogenous signaling in adult fly ovaries. Our results overall closely matched our genetic evidence of a negative regulation of Ras levels and MAPK signaling by RpS21. As in the eye, we found this negative regulation to preferentially impact weak Ras/MAPK signaling.

In the heads of *Ras^V12^Rare* flies, both transgenic Ras protein and MAPK signaling levels increase in *RpS21/+* animals relative to wild type. However, unlike our lack of an observable phenotypic enhancement of *Ras^V12^Common* in the eye, at the biochemical output level we also observe an increase in the level of Ras^V12^Common and MAPK signaling in the *RpS21*^0375^*/+* background (Fig 4b, **FigS3b**). This result shows that RpS21 also modifies Ras^V12^Common protein levels, but that only modification of Ras^V12^Rare protein by RpS21 leads to an observable phenotypic output. Next, we examined S2 cells transduced with an expression vector encoding either *Ras^V12^Common* versus *Ras^V12^Rare* and used RNAi to reduce RpS21 levels. Consistent with our mutant analysis in animals, RNAi reduction of RpS21 also elevates Ras^V12^Rare levels and MAPK signaling in *Ras^V12^Rare* cells. As in our fly eye analysis, we did not observe an effect of RpS21 reduction on Ras/MAPK pathway protein levels or activation in *Ras^V12^Common* cells. We note that relative to *Ras^V12^Rare* S2 cells, the level of Ras^V12^ protein and MAPK signaling are substantially elevated independently of RpS21 knockdown in *Ras^V12^Common* cells (Fig 4c, **FigS3c**), perhaps suggesting that Ras signaling is above a threshold of modification in this context. These findings in the adult head and in S2 cells imply that RpS21 regulates Ras/MAPK signaling in a manner that depends on signal strength.

Although the above results show that codon-altered *Ras* transgenes can be used to experimentally manipulate signal strength, the ability of RpS21 to regulate Ras/MAPK should extend beyond codon content, and therefore should impact endogenous MAPK signaling. We gained support of the idea that weaker, endogenous Ras/MAPK signaling is susceptible to *RpS21* knockdown irrespective of codons, through our examination of another cell type, cultured *Drosophila* KC cells. In these cells, irrespective of transgenic MAPK activation, p-Erk is increased upon addition of serum following a period of serum starvation. We find that upon re-addition of serum (*see* Methods), *RpS21 RNAi* elevates endogenous p-Erk levels (**FigS3d**), again consistent with a negative regulation of the pathway by RpS21. To assess whether endogenous MAPK signaling can be regulated by RpS21 *in vivo*, we assessed the effect of disrupting one allele of the *RpS21* gene on endogenous MAPK signaling in the ovaries of flies, a tissue where EGFR/Erk signaling has a well-defined role^40, 41^ and where phosphorylated Mek and Erk are readily detected (Fig 4d). In this tissue, endogenous p-Mek and p-Erk levels increase in both *Df(2L)*BSC692/+ and *RpS21^0375^/+* animals relative to control *w^1118^* animals (Fig 4d). Together, these findings demonstrate that RpS21 negatively regulates endogenous Ras/MAPK signaling in two contexts (KC cells and adult ovaries).

Our immunoblot analysis validates our genetic screen finding that RpS21 can act to negatively regulate MAPK signaling, in a manner that potentially depends on the strength of Ras/MAPK signaling. One interpretation of these data is that RpS21 has a minimal effect on MAPK signaling output above a certain threshold of MAPK signaling. Such a model would predict that experimentally reducing the amount of *Ras^V12^Common* expression should render fly eye development sensitive to the *RpS21^0375^/+* mutant background. To experimentally test this threshold model, we took advantage of the well-known fact that expression of transgenes using the Gal4-UAS system is responsive to temperature, with higher temperature resulting in higher expression over the physiological range of 18°C-29°C. In agreement, increasing temperature from 18°C to 29°C causes an observable and significant increase in the phenotypic severity of *Ras^V12^Common* (and *Ras^V12^Native*), but has little effect on *Ras^V12^Rare* (**FigS3e**). These results showed that we could potentially use temperature to test a signaling threshold model. We thus evaluated the rough-eye phenotype of *sev-Ras^V12^Common* versus *sev-Ras^V12^Rare* flies in a wild-type versus *RpS21^0375^/+* mutant background, only this time at 18°C. At this lower temperature, *RpS21/+* now acts as an enhancer of *Ras^V12^Common* (Fig 4e). Interestingly, *RpS21/+* no longer enhances *Ras^V12^Rare,* underscoring the sensitivity of *RpS21/+* to Ras/MAPK signaling strength. Collectively, these results demonstrate that while RpS21 negatively regulates Ras-MAPK signaling in diverse contexts, at the phenotypic level this regulation preferentially impacts low level Ras/MAPK signaling.

### MAPK signaling can promote RpS21 expression

The MAPK pathway is known to control expression of its negative regulators, which can provide pathway feedback regualtion^42–44^. Further, *Drosophila* Ras/MAPK signaling was recently shown to regulate rRNA synthesis and ribosome biogenesis^45^. To investigate whether RpS21 can be similarly regulated by the MAPK pathway, we compared the level of RpS21 protein when this pathway was activated in S2 cells by expression of activated Mek^EE^, Ras^V12^, or Raf^ED^. All three forms of activating the MAPK pathway elevate levels of RpS21 (Fig 5a,b, **FigS4a,b**). We validated these results in the opposite direction, namely demonstrating that the high levels of RpS21 induced by *Ras^V12^Common* or *Ras^V12^Rare* in KC cells is reduced by treatment with the Mek inhibitor Trametinib (Fig 5c**, FigS4c**). *In vivo*, RpS21 levels are higher in heads of *sev-Ras^V12^Common* compared to *sev-Ras^V12^Rare* flies (Fig 5d**, FigS3d**), again consistent with a positive relationship between MAPK activation and RpS21 levels.

**Figure 5.**
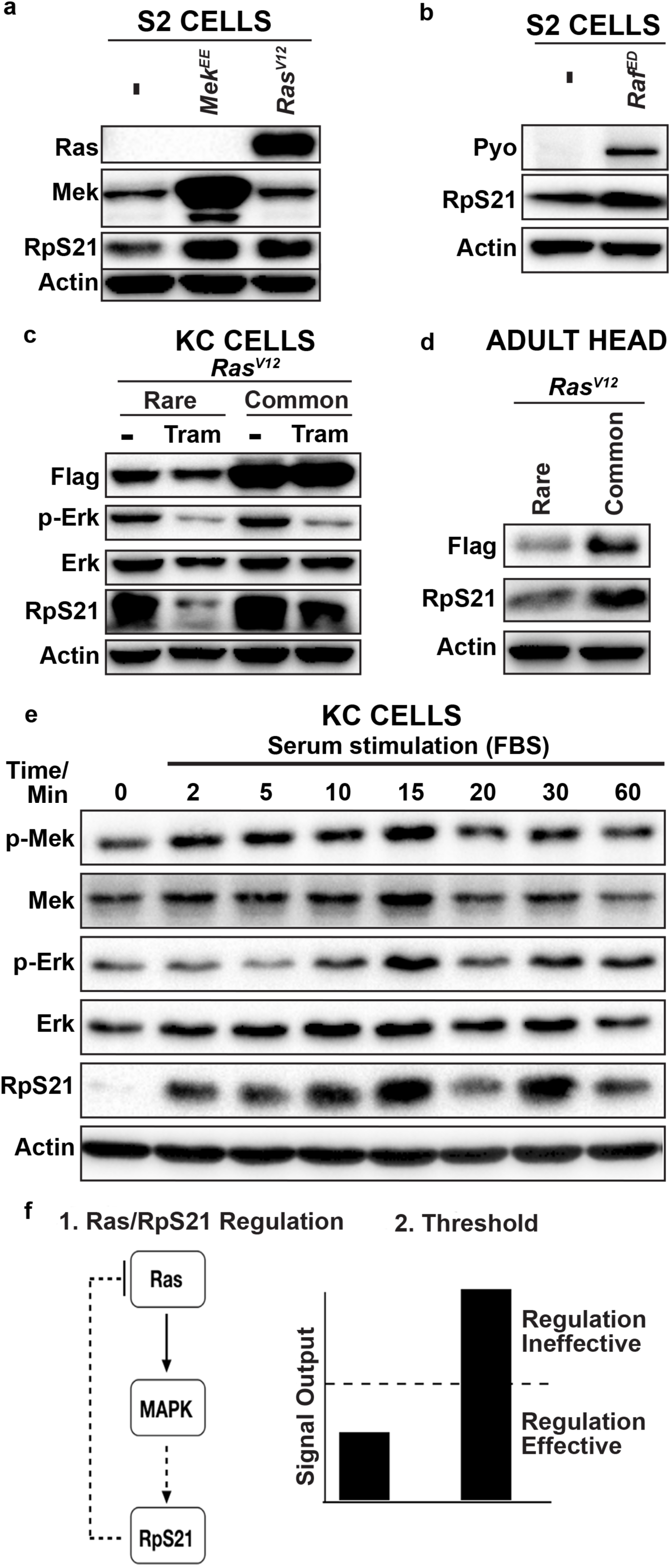
MAPK signaling can promote RpS21 expression. **(a,b)** Immunoblot detection of Ras, Mek, RpS21, Raf, or actin as a loading control in S2 cells stably transduced with expression vectors expressing **(a)** no transgene (-), activated Mek^EE^, or activated Ras^V12^ or **(b)** no transgene (-) or activated Raf^DD^ (anti-pyo antibody to detect tagged pyo-Raf^ED^). **(c)** Immunoblot detection of transgenic Ras^V12^ (with an anti-FLAG antibody), phosphorylated (p-) and total Erk, RpS21, and actin as a loading control from lysates derived from KC cells stably transduced with expression vectors expressing the indicated *Ras^V12^*transgenes in the absence (-) and presence of the Mek inhibitor trametinib (Tram). **(d**) Immunoblot detection of transgenic Ras^V12^ (with an anti-FLAG antibody), RpS21, and actin as a loading control from lysates derived from the head of flies with the indicated versions of transgenic *Ras^V12^*. **(e)** Immunoblot detection of phosphorylated (p-) and total Mek and Erk, RpS21, and actin as a loading control from lysates derived from serum-starved KC cells at the indicated times after stimulation with fetal bovine serum (FBS). **(f)** Proposed model of the effect of RpS21 on different levels of MAPK signaling.

The above results suggest that, as Ras/MAPK activation rises, so will RpS21 levels, which will then feedback to inhibit Ras/MAPK activation. To test this idea, we took advantage of our serum re-addition model in KC cells to explore the relationship of endogenous MAPK signaling and RpS21 levels in a temporal manner. We thus examined the amount of RpS21 expressed by immunoblot analysis when the MAPK pathway was stimulated in serum-starved KC cells by the addition of serum, as above. As expected, addition of serum causes a rapid increase in p-Mek followed by p-Erk levels, which is mirrored by a rapid increase in RpS21 levels (Fig 5e**, FigS4e**). These results are consistent with our findings that the MAPK pathway stimulates RpS21 expression. Interestingly, after RpS21 levels reach their peak, there is a reduction in p-Mek and p-Erk (Fig 5e**, FigS4e**). This result is again consistent with RpS21 acting as a negative regulator of MAPK signaling. We note here that the timing of when RpS21 reached peak expression after serum stimulation varied from experiment to experiment and was influenced by cell type, but typically tracked with p-Mek and p-Erk levels (not shown). At later time points, both RpS21 and MAPK activity levels again rise (Fig 5e**, FigS4e**), which may reflect the known ability of MAPK signaling to oscillate^17, 46^. Collectively, these data support a model whereby RpS21 acts as both a negative regulator and effector of the MAPK pathway. However, as revealed in our genetic screen, this regulatory relationship preferentially manifests itself at low signal output levels (Fig 5f).

## DISCUSSION

Here, we revisit a well-proven strategy to identify Ras/MAPK modifiers (a haploinsufficiency screen in the *Drosophila* eye), but do so with the new angle of altering codon usage in a core signaling component to find signal strength-dependent regulators. We show here that changing codon usage in a signaling pathway component can be an effective strategy to find signal strength-dependent modifiers, as evidenced by our identification of 15 *Df* from a whole-genome screen that only modify the rough-eye phenotype driven by either a common or rare codon-enriched *Ras^V12^*transgene, but not both. From these efforts, we identify the *RpS21* gene as a negative regulator of a weak or low-level *Ras* phenotype in the *in vivo* context of eye development. We also provide evidence that the MAPK pathway can increase RpS21 expression, suggesting a potential negative feedback relationship between Ras/MAPK signaling and RpS21.

Our results show that altering codon usage can serve as a valuable platform to stably alter protein production to undertake signal strength-specific screens. Clearly, there are other ways that one can modulate signal output strength, such as modulating gene expression strength as we also do here. However, an advantage of altered codon usage is that it can be hard-wired into the genome, and thus no additional (and potentially confounding) experimental parameters such as altering temperature, inducing genes with drugs, etc. are required. Our approach should be applicable to any signal transduction pathway. The utility of our approach is underscored in the fact that signal strength-specific modifiers found in our screen appear to be enriched for genome regions not previously linked to *Ras* genetic modification. The causative genes contained within 14 of these differential *Df* hits remain to be mapped, and represent a potentially rich source of new genes modulating Ras/MAPK signaling. Intriguingly, these differential hits appear to be enriched in bHLH transcription factors. Of note, the bHLH transcription factor Myc is a well-known Erk target^47–50^, and it will be interesting to explore whether specific bHLH transcription factors are preferentially targeted by this pathway in signal strength-dependent contexts.

Given the importance of Ras/MAPK signaling in many settings across evolution, our identified modifiers may shed insight into how this pathway is controlled at different signal strengths. While our focus here is on *Drosophila* eye development, signal strength dependencies of the Ras/MAPK pathway are appreciated to play a role in human disease. Activating mutations in the MAPK pathway of humans underlie a class of human diseases termed RASopathies^51^. Further, relevant to our approach here, the weakest expressed mammalian RAS isoform, *KRAS,* is enriched in rare codons^23^ and is the most commonly mutated RAS isoform in human cancers^52^. As such, modifiers of weak Ras signaling may provide a new class of proteins to explore in these diseases.

Our approach found that RpS21 functions as a negative regulator of low Ras/MAPK signaling. As Ras/MAPK signaling is known to drive tissue growth in diverse settings, this may suggest that RpS21 can function as a negative regulator of tissue or tumor growth. Interestingly, downregulation of RpS21 was previously shown to cause excessive hyperplasia in hematopoietic organs and imaginal disc overgrowth during larval development, suggesting RpS21 acts as tumor suppressor in *Drosophila*^39^. Although this finding may seem paradoxical given that ribosomal mutants in flies are well-known to cause “minute” phenotypes, characterized by short bristles, small body size, and delayed growth^53–56^, a subset of ribosomal proteins including RpS21 have been identified to have a growth suppressive role^39, 57–61^, and our ribosomal candidate screen may have identified other such subunits. Further, haploinsufficiency of many ribosomal proteins is reported to be tumorigenic in zebrafish^62^, and heterozygous inactivating mutations of ribosomal proteins have been described in human cancers^63, 64^. Several mechanisms have been proposed to account for this apparent tumor suppressor activity of ribosome protein downregulation, including activation of p53^65–67^, inhibition of NF-KB^68^, E2F^69^, MYC^70^, and CDK8^71^. Thus, RpS21 joins the ranks of an emerging number of ribosomal proteins with roles in growth suppression, although whether RpS21 acts as a tumor suppressor in mammals awaits investigation.

The mechanism underlying the negative regulation of Ras/MAPK signaling by RpS21 remains to be determined. In our work, we found that RpS21 downregulation promotes elevated levels of Ras^V12^ protein. The effect of RpS21 on Ras^V12^ protein level could potentially be through RpS21’s canonical ribosomal function or through an extra-ribosomal function. Dose-dependent ribosome dysfunction is linked to the human disease Diamond-Blackfan anemia, where heterozygous mutations in specific ribosomal subunits are linked, at least in part, to compromised ribosome biogenesis and translation^72–75^. A defect in RpS21 ribosomal function may trigger ribosomal biogenesis defects that alter translational fidelity or promote generation of oncoribosomes to preferentially express subset of mRNA pools^76, 77^. Alternatively, RpS21 might participate in other cellular processes independent of its canonical ribosomal function, as has been shown for other ribosomal subunits^78–81^. In contrast to our screen results revealing negative regulation by RpS21 in four different contexts, numerous ribosomal proteins (RpS21 included) were found among 1,162 genes to positively regulate Erk phosphorylation in a previous primary screen in cultured *Drosophila* S2R+ cells^14^. Unlike this Erk activation screen, we note that our *Ras^V12^* eye modifier screen hits were not preferentially enriched for ribosomal subunits, and that ribosomes in general are not enriched among known FlyBase Ras genetic enhancers/suppressors. We hypothesize that the addition of insulin to the growth media, required for Erk activation in the context of the S2R+ cell screen, revealed a dependency for cell growth, which is dependent on both ribosomes and Erk activation. S2R+ cells have known differences from S2 cells in response to external signaling, and this could reflect differences in MAPK regulation in this context as well^82^, underscoring the need to understand signaling dynamics and regulation in a given biological context.

Finally, our work may have uncovered another layer in the increasingly appreciated negative feedback regulation of Ras/MAPK signaling^44^, one that preferentially operates at a low signal output threshold. Many negative regulators of Ras/MAPK signaling are involved in feedback loops and are conserved in *Drosophila*, *C elegans*, and humans^9,43, 83–85^. In this regard, we present evidence that RpS21 not only suppresses Ras/MAPK signaling, but that Ras/MAPK signaling can stimulate RpS21 production. Future work can determine whether this elevation of RpS21 is a result of increased free (non-ribosomal) RpS21, or ribosomes in general. Nevertheless, our data may suggest the intriguing possibility that RpS21 is a negative regulator of the Ras/MAPK pathway as part of a negative feedback loop. Related to this, another question for future investigation is why RpS21/MAPK co-regulation is non-functional in contexts of heightened Ras/MAPK signaling, as we observed in S2 cells with high Ras/MAPK biochemical output, as well as at the phenotypic output level where *Rps21/+* failed to noticeably modify the eye phenotype of *Ras^V12^Common*. One possible explanation is that different MAPK signaling strengths activate a different host of MAPK targets, and this impacts the degree of negative regulation by RpS21. To that end, it will be important to further mine our screen to identify single gene modifiers in the other 14 *Dfs*, which may similarly yield new regulatory insight into the Ras/MAPK pathway.

## METHODS

### Generation of codon-altered Ras^V12^ genes in Drosophila

Codon-altered exon sequences for *Ras^V12^Common* and *Ras^V12^Rare* were created using the Kazusa codon usage database (https://www.kazusa.or.jp/codon/) and subsequently generated by Invitrogen GeneArt Gene Synthesis (ThermoFisher Scientific). A cDNA clone (LD17536, Drosophila Genomics Resource Center) was used as a template to generate the non-altered *Ras85D* sequence. To generate *Ras^V12^Native*, the QuikChange II Site-Directed Mutagenesis Kit (Agilent) was used to change codon 12 in *Ras85D* from GGA (glycine) to GTA (valine). Subsequently, primers (sequences available upon request) were designed to amplify *Ras* sequences and the Invitrogen Gateway BP Clonase II Enzyme Mix (ThermoFisher Scientific) was used to insert these sequences into the Gateway entry vector pDONOR221 (ThermoFisher Scientific). Subsequently, the Invitrogen LR Clonase Enzyme Mix (ThermoFisher Scientific) was used to insert the *Ras^WT^*, and *Ras^V12^* Native, Common, and Rare sequences into the Gateway destination vector pBID-UASC-FG (Addgene Plasmid #3520^86^), which has a N-terminal FLAG tag and a PhiC31 site for site-directed genomic insertion. pBID-UASC-FG-*Ras* plasmids were prepared with a ZymoPURE II Plasmid Midiprep Kit (Zymo Research) and sent to Model System Injections (Durham, NC, USA) for injection into *attP40 (2L)* flies. For cell culture, *Ras^V12^Common* and *Ras^V12^Rare* transgenes were cloned into pMKInt-Hyg vectors, which were sequenced to confirm the correct sequence.

### Fly stocks

All flies were raised at 25°C on standard media unless noted otherwise (Archon Scientific, Durham NC). FlyBase (http://FlyBase.org) describes full genotypes for all stocks used in this study. See TableS1 for Df stock information and TableS2 for ribosomal mutant stock information (except *RpS21).* All other stocks were the following genotypes (Bloomington Drosophila Stock Center numbers in parentheses): *ksr^S−627^/TM3,Sb* (#5683), *aop^yan-XE18^/CyO* (#8777), *betaggt-I^S-2554^* (#5681), *RpS21^03575^/CyO* (#11339), and the Bloomington Deficiency(Df) kit. The following stocks were generated for this study: *UAS-FLAG-Ras*, *UAS-FLAG-Ras^V12^Native*, *UAS-FLAG- Ras^V12^Common*, and *UAS-FLAG-Ras^V12^Rare*.

### Fly Genetics and Deficiency Screen

The *Ras* transgenes were combined with a *sev*Gal4 driver and subsequently crossed to *Df*/Balancer flies. After 18-20 days, the rough eye phenotype of the resulting progeny was scored. The scoring system was as follows (<u>category=numerical score</u>, qualitative description): <u>Mild=1</u>, no discoloration or necrotic tissue; <u>Moderate=3</u>, discoloration and no necrotic tissue; <u>Severe=5</u>, discoloration and necrotic tissue (see Fig 1b,c). Severity scores for each genotype was calculated as follows: (#Mildx1+#Moderatex3+#Severex5)/Total # of flies. To determine if haploinsufficiency for a subset of genes altered the rough eye phenotype the following two genotypes for each deficiency (*Df*) were compared: *Ras* transgene only and *Ras* transgene + *Df* (used as an internal comparison to control for background effects). Then, we calculated a fold change score for both *Ras^V12^Common* and *Ras^V12^Rare* for each deficiency: *Ras* transgene + deficiency/*Ras* transgene. For the primary screen, the fold change score was defined as follows: enhancer (fold change ≥1.35 or 5X more flies eclosed); suppressor (fold change ≤0.65 or 5X less flies eclosed). For the secondary screen, the fold change score was defined as follows: enhancer (fold change ≥1.95 or 5X more flies eclosed); suppressor (fold change ≤0.50 or 5X less flies eclosed). The final phenotype for a deficiency was defined as follows: <u>not a modifier</u> (neither *Ras^V12^Common* or *Ras^V12^Rare + Df* were modified); <u>enhancer</u> (both *Ras^V12^Common* and *Ras^V12^Rare + Df* were enhanced); <u>suppressor</u> (both *Ras^V12^Common* and *Ras^V12^Rare + Df* were enhanced); <u>differential</u> (only *Ras^V12^Common* or *Ras^V12^Rare + Df* were modified). Images of fly eyes were obtained using a Leica MZ10F microscope with a PlanApo 1.6X objective, Pixel Shift Camera DMC6200, and LASX software.

### Protein Preparation and Analysis

All protein samples were prepared by homogenizing tissue on ice. For Fig 1d, samples were processed in Laemmli buffer and then boiled for 5min. Samples were separated by 12% sodium dodecyl sulfate-polyacrylamide electrophoresis (SDS-PAGE) gels and transferred to an Odyssey nitrocellulose membrane (LI-COR Biosciences) for immunoblotting. The following antibodies were used: anti-FLAG M2 (1:500, Sigma, anti-mouse), anti-α-tubulin (1:20,000, Sigma, anti-mouse), IRDye 800CW (1:20,000, LI-COR Biosciences, anti-mouse). Signal was detected using LI-COR Odyssey CLx and analyzed using Image Studio (LI-COR Biosciences). For all other immunoblots, samples were processed in RIPA buffer containing 1% IGEPAL, 50mM NaCl, 2mM EDTA, 100mM Tris-HCl, pH 8.0, 0.1% Glycerol, 50 mM Naf, 10mM Na3VO4, and protease inhibitors (Roche). *Drosophila* heads and ovaries were collected and transferred to cold lyses to be homogenized with a pellet pestle. Lysates were incubated at 4 °C for 30 min on end-to-end rotator and then centrifuged at 21,000×g for 10 min. The supernatant was transferred to a new tube. Total protein was quantified using a BCA kit (Bio-Rad) and either 30 or 50 micrograms of protein was used for separation on either 12.5% or 15% gradient SDS-PAGE gels. Proteins on SDS gels were transferred onto polyvinylidene difluoride membranes. These membranes were probed with anti-Flag (Sigma, anti-mouse 1:1000), anti-Pan-Ras (Millipore, 1:50), anti-β-actin (Cell Signaling, 1:1000), anti-p-MEK1/2 (Cell Signaling, 1:500), anti-MEK1/2 (Cell Signaling, 1;500), anti-p- ERK1/2 (Cell Signaling, 1:1000), anti-ERK1/2 (Cell Signaling, 1:1000), anti-RpS21 (Abcam, 1:2000), anti-Pyo (anti-Glu-Glu, VWR, 1:500) primary antibodies in blocking buffer containing 5% milk followed by the secondary antibodies of goat anti-mouse IgG (H+L) HRP (Life Technologies, 1:10000) or goat anti-rabbit IgG (H+L) HRP (Thermo Fisher Scientific, 1:10000). Immunoblots were visualized using Clarity Max™ ECL Western Blotting Detection Reagent (Bio-Rad) followed by exposure to digital acquisition using Chemi Doc Imager (Bio-Rad). For all blots, the contrast and/or brightness were altered equally across the entire image and then images were cropped for displaying as figures.

### Cell Culture

KC and S2 cell lines were obtained from Bloomington (Indiana University DGRC Bloomington) and as a gift from Dr. David MacAlpine (Duke University) respectively. These cells were cultured in Schneider’s *Drosophila* medium (Invitrogen) supplemented with 10% fetal bovine serum (FBS) and 1% penicillin–streptomycin-L Glutamine (Invitrogen) at 25°C. FBS was heated for 60 minutes in 58°C and then cool down before added to medium. These cells were confirmed to be free of mycoplasma infection, as measured by the Duke Cell Culture Facility using MycoAlert PLUS test (Lonza). S2 and KC cell lines were stably transduced with the pMKInt-Hyg vector encoding *Ras^V12^Common* and *Ras^V12^Rare* cDNAs using 1000 ng of DNA in 6 well plates as instructed by Effectene transfection reagent (Qiagen). The following day, Schneider’s media was changed, and cells were seeded in a coated culture dish (100×20 mm). Four days later, cells were passaged with fresh Schneider’s medium and 200 µg/ ml hygromycin (Invitrogen) was added. The stably transfected cells were selected within a month growing in media containing hygromycin. Three days prior to any experiment, these cells were grown in media without hygromycin. Four million S2 cells that were stably transduced with *Ras^V12^Common* and *Ras^V12^Rare* plasmids were seeded into coated tissue culture dishes (60×15mm, VWR) with 2 ml of Schneider’s media (without FBS). Sixty micrograms of RpS21 dsRNA were added on top of these cells. One hour later, two ml Schneider’s media containing 20% FBS were added on top of 2 ml Schneider’s media without FBS resulting in medium with 10% FBS concentration in total media of this culture. Within 16-24 hours after RNAi treatment, expression of *Ras^V12^Common* and *Ras^V12^Rare* transgenes were induced by CuSO4 for another 12 hours. Finally, these cells were collected 30-36 hours after dsRNA treatment. S2 cells were transiently transfected with either pMet-HA-Ras^V12^, pMet-pyo-Raf^ED^, and pMet-myc-Mek^EE^ vectors (kindly provided by Dr. Marc Therrien^15^) using the Effectene transfection reagent as described above. Two days later, these cells were treated with 500 µM CuSO4 for 16 hours to include expression of transgene. KC cells *Ras^V12^Common* and *Ras^V12^Rare* were treated with 10 nM trametinib for 12- 16 hours prior collecting for immunoblot assay.

### dsRNA synthesis

S2 cell DNA was used to produce a PCR template for RpS21 dsRNA production using the forward primer “TAATACGACTCACTATAGGGTTACTGACCAGCCGATACCC” and reverse primer “TAATACGACTCACTATAGGGCCACGCTTAGAAGTTCCTGC”. Next, 500 ng RpS21 PCR template was used for an *in vitro* production of dsRNA as instructed in the MEGAscrip T7 transcription kit (ThermoFisher). The dsRNA solution was cleared using MegaClear™ kit (ThermoFisher). Finally, the concentration of *RpS21* dsRNA was measured and stored in −80°C for future use.

### Gene enrichment analysis and statistical analyses

To determine the Codon Adaptive Index (CAI), sequences were entered at the CAIcal web-server (http://genomes.urv.es/CAIcal^87^. For gene enrichment, deficiency sequence boundaries were defined using coordinates available through FlyBase^88^ and the Bloomington *Drosophila* Stock Center website. Deficiencies were then uploaded as a custom BED track to the UCSC Genome Browser (Reference Assembly ID: dm6). Genes overlapping the deficiency coordinates were then extracted using BEDtools for additional analysis^89^. A deficiency was determined to contain known *Ras* modifiers if any of the deficiency covered genes known as *Drosophila Ras85D* genetic interactors (332 interactors, FlyBase). Enhancers and suppressor deficiencies were analyzed using the same metric against known *Ras85D* interactors of the same respective modifier type. Statistical analysis (chi-square) was performed using Graphpad Prism v8.1. FlyBase Gene Group Enrichment analysis was performed by comparing deficiency covered genes with pre-defined FlyBase Gene Groups. Analysis and statistical tests were performed in R using Gene Overlap package (https://rdrr.io/bioc/GeneOverlap/) and results are reported as adjusted p-values (False Discovery Rate^90^, using Benjamini Hochberg correction). Graphs and statistical analyses were generated using GraphPad Prism 7. Statistical tests and P-values are detailed in figure legends. For all tests, P-value reporting is as follows: (P>0.05, n.s.; P<0.05,*; P<0.01,**; P<0.001,***, P<0.0001,****).

## ACKNOWLEDGMENTS

We thank David MacAlpine and Marc Therrien for providing useful reagents, Heather MacAlpine for technical help with *Drosophila* cell culture, and David MacAlpine, Bernard Mathey-Prevot, and Daniel Lew for helpful critiques on the manuscript. This project was supported by American Cancer Society research scholar grant RSG-128945 a Duke Cancer Institute Pilot project grant to DF, National Institutes of Health grants R01CA94184 and P01CA203657 to CC, and a National Institutes of Health grant GM109538 to JS.

## Supplementary Tables

**TableS1.** Raw data for the primary and secondary eye screens (attached separately).

**TableS2.** Tests of genetic interaction between *Ras^V12^* Rare/Common and small ribosomal subunits.

| Ribosome Gene | Allele/RNAi | Type of Modifier |
| --- | --- | --- |
| *RpS2* | *RpS2*<sup>PRW1</sup> | Enhancer of Rare |
|  | *RpS2*<sup>k01215</sup> | Not a modifier |
| *RpS3* | *RpS3*<sup>Plac92</sup> | Suppressor of both |
|  | *RpS3*<sup>2</sup> | Not a modifier |
| *RpS9* | *RpS9*<sup>f03376</sup> | Suppressor of both |
| *RpS12* | *RpS12*<sup>s2783</sup> | Not a modifier |
|  | *RpS12*<sup>EY23269</sup> | Not a modifier |
| *RpS13* | *RpS13*<sup>1</sup> | Not a modifier |
| *RpS15* | *RpS15*<sup>e01611</sup> | Enhancer of both |
|  | * *RpS15* <sup>HMS04362</sup> | Not a modifier |
| *RpS16* | * *RpS16* <sup>KK100610</sup> | Enhancer of Common |
| *RpS17* | *RpS17*<sup>6</sup> | Not a modifier |
| *RpS23* | *RpS23*<sup>KG07144</sup> | Enhancer of Rare |
| *RpS24* | *RpS24*<sup>f06717</sup> | Enhancer of both |
|  | *RpS24*<sup>SH2053</sup> | Enhancer of both |
| *RpS26* | *RpS26*<sup>04553</sup> | Not a modifier |
|  | * *RpS26* <sup>KK109004</sup> | Enhancer of both |
|  | * *RpS26* <sup>HMS00270</sup> | Not a modifier |
| *RpS27A* | *RpS27A*<sup>mfs31</sup> | Enhancer of Rare |
|  | *RpS27A*<sup>04820</sup> | Not a modifier |
| *RpS28A* | * *RpS28A* <sup>KK112278</sup> | Enhancer of both |
| *RpS28B* | * *RpS28B* <sup>HMS00635</sup> | Enhancer of both |
|  | * *RpS28B* <sup>KK109420</sup> | Not a modifier |
| *RpS29* | *RpS29*<sup>A074</sup> | Not a modifier |

\*=RNAi

## Supplementary Figure Legends

**Figure S1.**
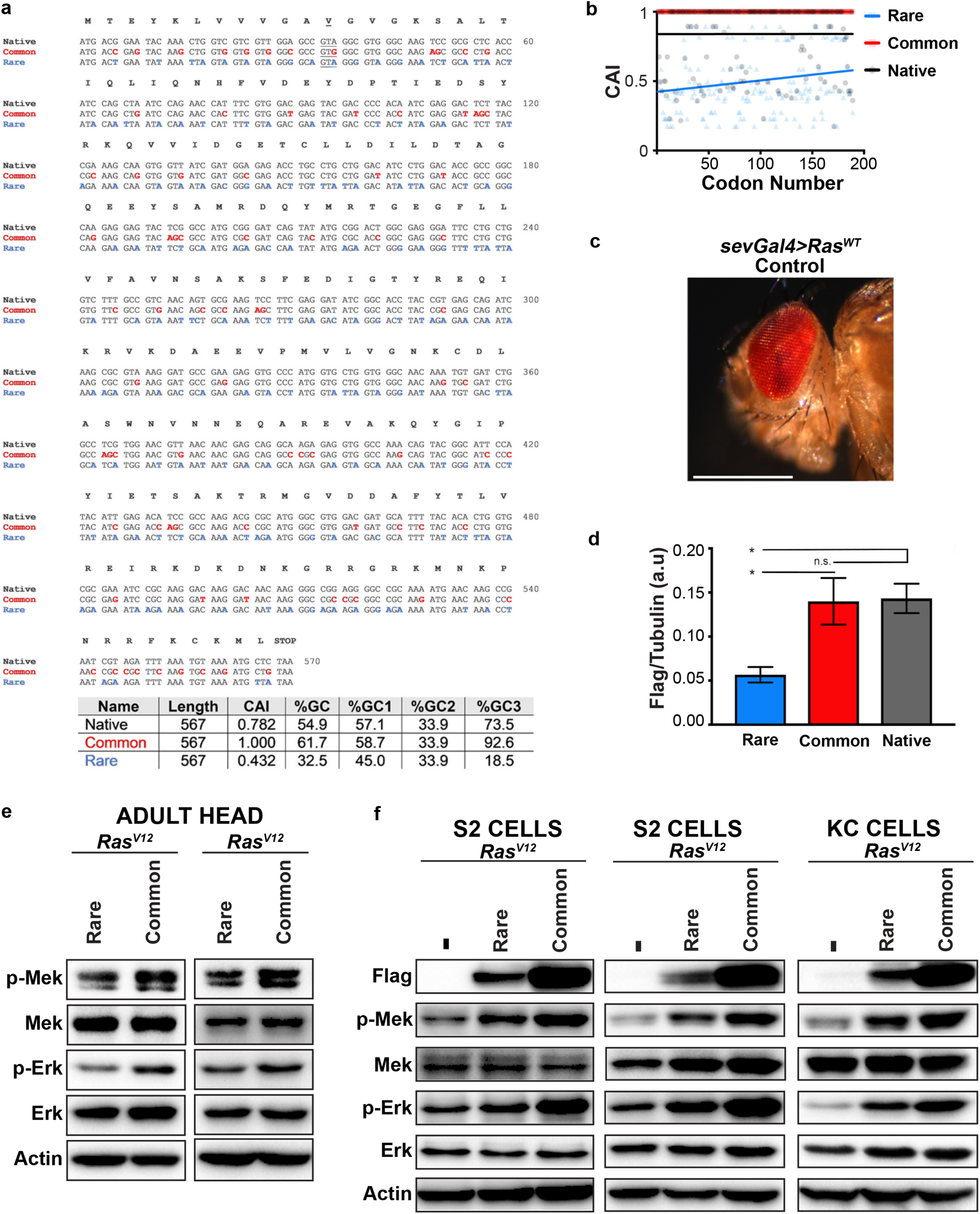
Codon manipulation of *Ras^V12^* promotes differential MAPK signal strength levels in *Drosophila*. **(a)** Alignments of *Ras* transgenes. Nucleotide changes highlighted for *Ras^V12^Common* (red) and *Ras^V12^Rare* (blue). Table with overall CAI score and GC content for *Ras* transgenes. (**b)** Codon Adaptive Index (CAI) plot. Transparent circles, squares, and triangles are individual CAIs per codon. Solid lines represent a best-fit line of individual points for each transgene. **(c)** Representative image of adult eye from animal expressing *sevGal4>Ras^WT^* **(d)** Quantification of protein levels at 25°C for blot in **Fig1d**. a.u.=arbitrary units. Data represent mean ± SEM, 3 replicates, Tukey’s multiple comparisons test. **(e)** Immunoblot detection of transgenic Ras^V12^ (with an anti-FLAG antibody), phosphorylated (p-) and total Mek and Erk, and actin as a loading control from lysates derived from **(e)** the head of flies with the indicated versions of transgenic *Ras^V12^*or **(f)** S2 cells stably transduced with expression vectors expressing the indicated *Ras^V12^* transgenes. First lane is S2 cells without any transfection.

**Figure S2.**
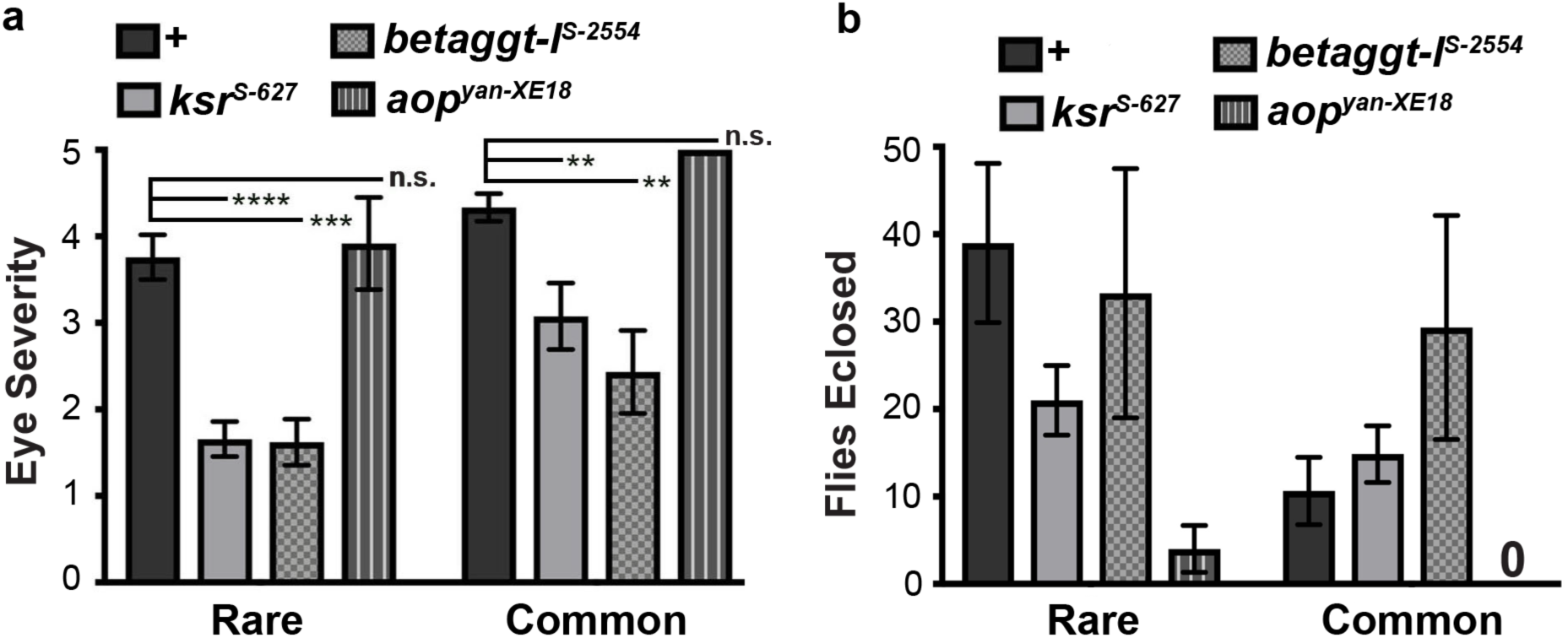
Known *Ras* modifiers alter phenotypes driven by codon-altered *Ras* transgenes. **(a)** Quantification of eye severity scores for *Ras* transgenes that are also haploinsufficient for known *Ras* modifiers. Data represent mean ± SEM, multiple replicates (using Dennett’s multiple comparison test). **(b)** The average number of flies eclosed per experiment for Rare and Common transgenes in a known *Ras* modifier background.

**Figure S3.**
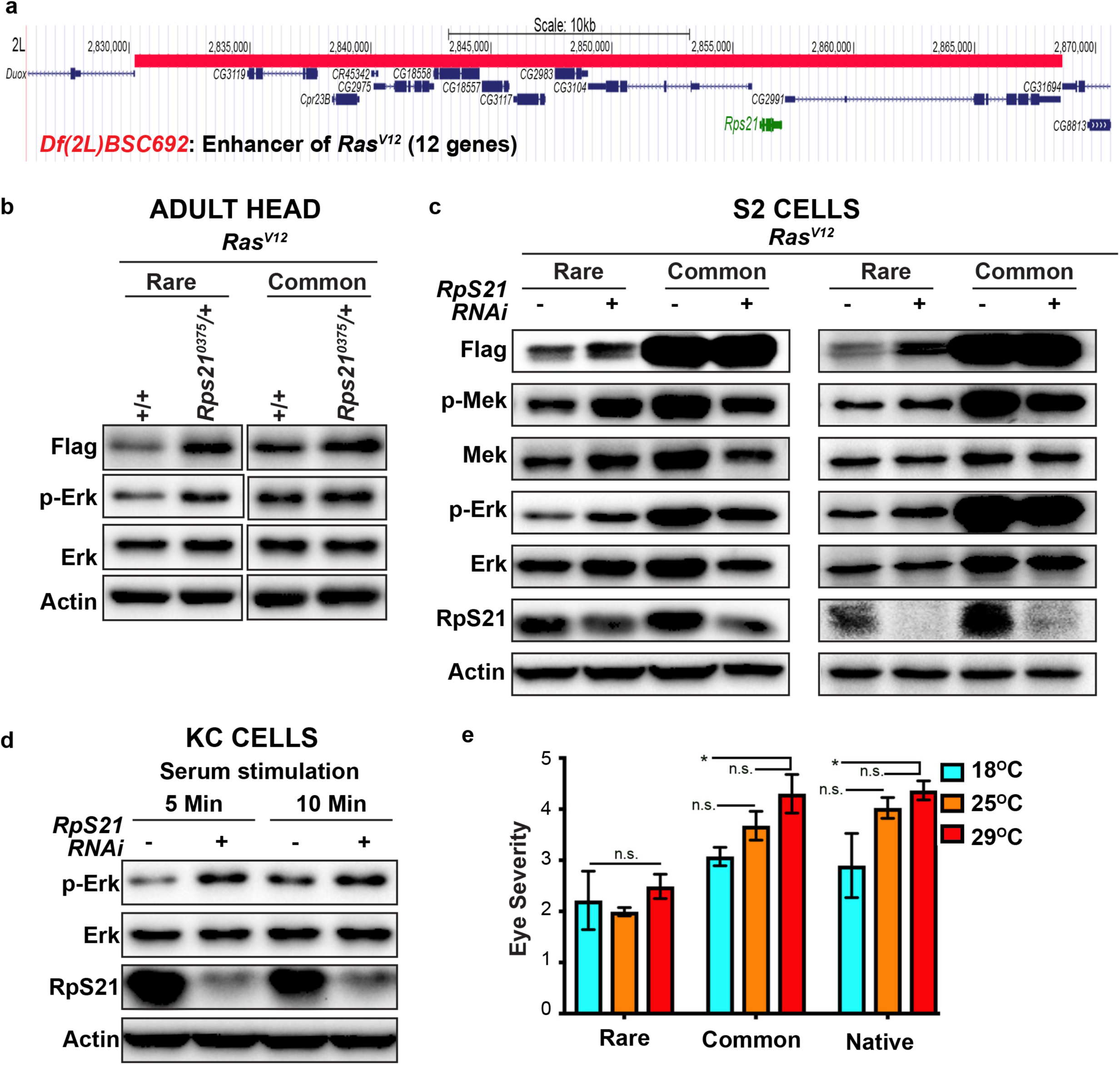
Rps21 negatively regulates Ras/MAPK signaling in settings of low signal output. **(a)** Genome map of *Df(2L)BSC692*. *RpS21* is highlighted in green. **(b, c, d)** Immunoblot detection of transgenic Ras^V12^ (with an anti-FLAG antibody), phosphorylated (p-) and total Mek and/or Erk, RpS21, and actin as a loading control from lysates derived from **(b)** the head of flies with the indicated versions of transgenic *Ras^V12^* in either the wild-type (+/+) or mutant (*RpS21^0375^*/+) RpS21 backgrounds, **(c)** S2 cells stably transduced with expression vectors expressing the indicated *Ras^V12^* transgenes in the absence (-) and presence (+) of *RpS21 RNAi* (Data represent two independent replicates.) or **(d)** KC cells stably transduced with expression vectors expressing indicated RasV12 transgenes in the absence (-) and presence (+) of *RpS21 RNAi*. **(e)** The mean ± SEM eye severity score of the genotypes from three replicate experiments at 18°C, 25°C, and 29°C. Tukey’s multiple comparisons test was used for statistical comparisons. \**p*<0.05. n.s., not significant.

**Figure S4.**
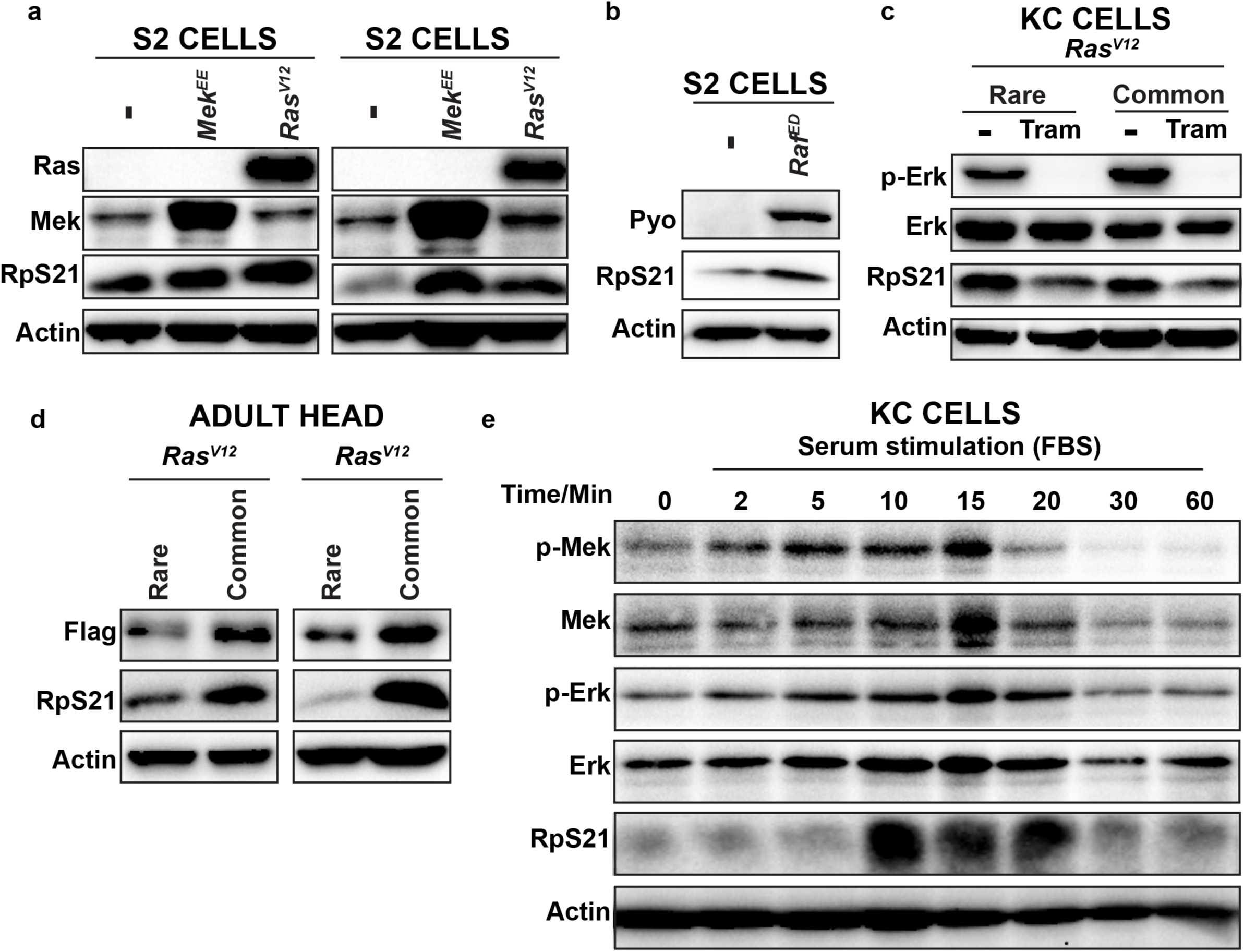
RpS21 expression is elevated by MAPK signaling. **(a,b)** Immunoblot detection of Ras, Mek, RpS21, Raf, or actin as a loading control in S2 cells stably transduced with expression vectors expressing **(a)** no transgene (-), activated *Mek^EE^*, or activated *Ras^V12^* (using anti-Pan Ras antibody to detect Ras^V12^) or **(b)** no transgene (-) or activated *Raf^ED^* (anti-Pyo antibody to detect tagged *pyo-Raf^ED^*). **(c)** Immunoblot detection of transgenic Ras^V12^ (with an anti-FLAG antibody), phosphorylated (p-) and total Erk, RpS21, and actin as a loading control from lysates derived from KC cells stably transduced with expression vectors expressing the indicated *Ras^V12^*transgenes in the absence (-) and presence of the Mek inhibitor trametinib (Tram). **(d)** Immunoblot detection of transgenic *Ras^V12^* (with an anti-FLAG antibody), RpS21, and actin as a loading control from lysates derived from the head of flies with the indicated versions of transgenic *Ras^V12^*. **(e)** Immunoblot detection of phosphorylated (p-) and total Mek and Erk, RpS21, and actin as a loading control from lysates derived from serum-starved KC cells at the indicated times after stimulation with fetal bovine serum (FBS).

